# Epstein-Barr virus BRRF1 induces butyrophilin 2A1 in nasopharyngeal carcinoma cells via the IL-22/JAK3-STAT3 pathway

**DOI:** 10.1101/2023.08.16.553646

**Authors:** Yue Liu, Zuodong Ye, Luo Chen, Allen Ka Loon Cheung

**Author notes:** Correspondence to: Dr. Allen KL Cheung, Rm 806, Run Run Shaw Building, Ho Sin-hang campus, Hong Kong Baptist University, 224 Waterloo Road, Kowloon Tong, Kowloon, Hong Kong SAR, China.

## Abstract

Epstein-Barr virus is highly associated with nasopharyngeal carcinoma (NPC) with genes expressed for tumor transformation or maintenance of viral latency, but there are certain genes that can modulate immune molecules. Butyrophilin 2A1 (BTN2A1) is an important activating protein for presenting phosphoantigen presentation for recognition by Vγ9Vδ2 T cells to achieve anti-tumor activities. We have previously shown that Vγ9Vδ2 T cells achieve efficacy against NPC when BTN2A1 and BTN3A1 are upregulated by stimulating EBV gene expression, particularly LMP1. Whilst BTN3A1 can be induced by LMP1 mediated IFN-γ/JNK/NLRC5 pathway, the viral gene that can regulate BTN2A1 remains elusive. Here, we showed that BTN2A1 expression is directly mediated by EBV BRRF1, which can trigger BTN2A1 promoter and downstream JAK3-STAT3 pathway in NPC43 cells as enlightened by RNA-seq data and verified by inhibitor experiments. Further, BRRF1 downregulated IL-22 binding protein (IL-22RA2) to complement the EBNA1-targeting probe (P_4_) induced IL-22 expression. These functions act in concordance to enhance the expression of BTN2A1, which likely lead to increased tumor cell killing by Vγ9Vδ2 T cells. Overall, this study elucidated a new mechanism of stimulating BTN2A1 expression in nasopharyngeal carcinoma cells by the EBV gene BRRF1.

**Summary statement:** This study demonstrated a new function Epstein-Barr Virus BRRF1 to induce BTN2A1 in nasopharyngeal carcinoma via the IL-22/JAK3-STAT3 pathway, which potentiates Vγ9Vδ2 T cells against NPC.

## Introduction

Nasopharyngeal carcinoma (NPC) is a type of malignant cancer that occurs in the nasopharynx. The etiological factor of NPC is not fully understood, but associations with Epstein-Barr Virus (EBV), alcohol, smoking and diet has been made, such as cured fish [1]. There is higher incidence of NPC in southern China regions, including Hong Kong and Taiwan, as well as Inguiny in Greenland [2]. Usually, diagnosis of NPC occurs at the late stages when symptoms become obvious, and classical surgery, chemotherapy and/or radiotherapy are not effective in eliminating the cancer [3]. Therefore, development of immunotherapy such as the use of immune cells has increasing potential for treating NPC patients when other options fail [4]. However, for the success of such new therapies, the understanding of the prevailing EBV within the NPC and whether its viral genes have any relationship with promoting anti-tumor immune responses is one avenue to explore.

Some latent EBV-encoded proteins, such as EBV-encoded nuclear antigen 1 (EBNA1), is persistently expressed and can promote tumorigenesis. EBNA1 is found in all EBV latently infected cells, and thus it can be used as a target for manipulation [5]. We, and others, have previously used EBNA1-targeting probes (L_2_)P_4_ and ZRL_5_P_4_, which emit fluorescent signals when they bind with the EBNA1 protein, to stimulate reactivation of the latent viral genome [6-8]. Specifically, we found that the (L_2_)P_4_ probe can trigger EBV LMP1 expression that promote the expression of butyrophilin (BTN), which are essential to activate the anti-tumor cytotoxic function of a type of innate immune T cells named γδ T cells. Besides latent membrane protein 1 (LMP1), other EBV genes could still have the possibility to be associated with BTN expression, such as BRRF1 as indicated by our previous analysis [9].

BRRF1 is not a well-characterized gene. Although it is encoded in the same region as the BZLF1 and BRLF1 EBV gene products. BRRF1 has been reported to enhance BRLF1-induced reactivation in gastric cancer cells and HEK293 cells [10]. Knockdown of endogenous BRRF1 inhibits chemical-induced reactivation in NPC cell lines [11]. BRRF1 was also shown as an inducer of cleaved protein expression [11]. It has been suggested that BRRF1 can interact with TRAF2 and mediate several major signaling pathways, including NF-κB and JNK activation [12]. In EBV-infected B cells, TRAF2 interacts with LMP1, which activates the NF-κB pathway [13]. In addition, studies have shown that BRRF1 and the p53 tumor-suppressing protein can work together to drive lytic gene expression within the epithelium [14]. BRRF1 can also act as a transcription factor by promoting the transcription function of c-Jun and activating the BZLF1 promoter through the cAMP Response Element (CRE) motif sequence [15]. From these studies, the function of BRRF1 has been attributed to mediating host cell pathways or other EBV genes that are related to EBV infection, however, there has been no indication that BRRF1 can affect the expression of BTN molecules that affect γδ T cells.

Vγ9Vδ2 T cells is a subtype of human T cells that has natural tumoricidal activities [16]. They are primarily activated in a T cell receptor (TCR)-dependent manner by the recognition of phosphoantigen (pAg) presented by the complex formed between BTN2A1 and BTN3A1 molecules [17, 18]. BTN are a family of transmembrane glycoproteins that play important roles in the modulation of cellular and immune responses [19]. The intracellular B30.2 domain of BTN3A1 is essential for recognizing the pAg as well as activating the Vγ9Vδ2-TCR [20], where its configuration and transport on the cell membrane is mediated by Rho GTPase [21]. Recent studies have shown that BTN2A1 binding to BTN3A1 is the rate-limiting factor and vital for Vγ9Vδ2 T cell cytotoxicity [22], where BTN2A1 binds to the Vγ9 chain of the TCR directly [17]. Thus, BTN2A1 and BTN3A1 molecules are particularly important for consideration in studying the effects of Vγ9Vδ2 T cells in immunotherapy against cancer. There have been several clinical trials that made use of these cells for immunotherapy against different cancers, including NPC [23], but the limited effectiveness is underexplored in the realm of BTN expression in the tumors. In this regard, we recently demonstrated that Vγ9Vδ2 T cells had enhanced efficacy against NPC when the EBNA1-targeting probe (P_4_) was used in conjunction in an *in vivo* mice model [9]. Further, we found that P_4_ resulted in the expression of LMP1, that lead to the activation of the IFN-γ/p-JNK/NLRC5 axis and induced a certain level of BTN2A1/BTN3A1 expression in the tumors [9]. Given that the expression of BTN2A1/BTN3A1 would be vital for the success of Vγ9Vδ2 T cell immunotherapy against NPC, we continue to explore if other EBV genes can directly stimulate BTN2A1 and BTN3A1 expression to achieve this.

In this study, we uncovered a new function of EBV BRRF1, where its expression can stimulate the expression of BTN2A1 in NPC cells. BRRF1 do so by triggering the promoter activity of BTN2A1 leading to increased mRNA and protein level of BTN2A1. In addition, we showed that while P_4_ could stimulate IL-22 cytokine expression in NPC43 cells, BRRF1 can downregulate the level of the inhibitory IL-22 binding protein. As a result, the IL-22 cytokine can activate the downstream JAK3-STAT3 pathway to achieve a greater increase in the expression of BTN2A1 in NPC.

## Materials and Methods

### Cell culture and plasmid

NPC cells (HK1, HK1-EBV, HONE1, and HONE1-EBV) were cultured using DMEM medium (GIBCO) supplemented with 10% fetal bovine serum (FBS; GIBCO). NPC cells derived from clinical specimens (C666-1 and NPC43) [24]were cultured in the RPMI-1640 medium (GIBCO) supplemented with 10% FBS. The culture medium for NPC43 was supplemented with ROCK inhibitor 4 μM Y27632 (Cat. No. S1049, Selleckchem). All cell culture media were supplemented 1% penicillin/streptomycin (GIBCO). The cells were maintained at 37°C and 5% CO_2_ incubator.

The expression plasmid encoding BRRF1 contains a Myc tag synthesized by the IGE company. The vector pcDNA3.1 was used as a control in the experiments.

### Isolation of total RNA and Reverse transcription quantitative real-time PCR (RT-qPCR)

RNAiso PLUS (Cat. No. 9109, Takara) was used to isolate total RNA from the cells. cDNA synthesis from total RNA using the Prime Script RT Reagent Kit (Cat. No. RR047A, Takara) according to the manufacturer’s instructions. cDNA was then used for qPCR analysis using the TB Green® Premix Ex Taq™ II (Ti RNase H Plus) (Cat. No. RR820A, Takara) according to the manufacturer’s instructions. GAPDH gene was used as housekeeping control, which was used for normalization in data analysis. **Table S1** shows the gene primer sets used in the qPCR reactions.

### RNA sequencing

Total RNA samples were sent to Novogene company to perform RNA sequencing based on the using the NovaSeq PE 150 platform. Three independent experimental samples from control and BRRF1-overexpressed cells were used for RNA sequencing analysis. 12 G of raw data per sample was generated for standard bioinformatic analysis.

### Cell transfection

NPC43 cells were seeded the day before transfection. Cell culture media was changed to Opti-MEM media (Cat. No. 31985062, GIBCO) 1 h before transfection. The transfectant plasmid DNA and PEI reagent was prepared in Opti-MEM. 3 μl of linear PEI (1mg/ml, Cat. No. 23966-1, Polysciences) transfection reagent was used per 1 μg of plasmid DNA. The transfectant mixture was gently vortexed, and left at room temperature for 20 min to allow DNA/PEI complexes to form. The control empty vector with PEI was used as vehicle control. The transfectant mixture was added to the cells uniformly and media changed to cell culture media after 5-6 h. The cells were incubated at 37°C and 5% CO_2_ before used for experiments.

### Inhibitor experiments

JAK3 signalling was inhibited by ritlecitinib (PF-06651600, Cat. No.: HY-100754, MedChemExpress) in the cell experiments. DMSO was used as control. Cells for experiments were seeded overnight to achieve 80% confluence before treated by ritlecitinib for one hour, followed by BRRF1 transfection, and incubated for another 24 hours before assessed.

### Western blotting analysis

Protein extracts from cells were assessed for protein concentrations using the Pierce BCA protein assay kit (Thermo Fisher Scientific). Equal amounts of protein lysates were added to SDS-PAGE gels and electrophoresis was performed, then transferred to the PVDF (Millipore) membrane using wet transfer system. The membranes were incubated with 5% Blocking-grade blocker non-fat milk (Cat no. 1706404, Bio-Rad) plus 0.5% Bovine Serum Albumin (BSA; Cat. No. A2153, Sigma) in Tris-buffered saline with 0.1% Tween-20 (TBS-T) for one hour at room temperature. Appropriate primary and secondary antibodies were then used for the membranes (**Table S2**), which may be cut into strips according to the protein of interest. Signals were visualized using the Pierce ECL Western Blotting Substrate (Cat. No. 32106, Thermo Fisher Scientific), and images were taken with ChemiDoc (Bio-Rad) and band intensities analyzed using ImageJ.

### Luciferase reporter assay

The BTN2A1, BTN3A1, and NLRC5 promoter regions were inserted into the pGL3 base plasmid to create luciferase reporter plasmids. As a negative control, the pGL3 plasmid vector was used. Sequencing was used to verify all constructs. The cytokine IL22 (10 ng/ml, Cat. No. 13059-HNAE, Sino Biological) was used in certain experiments after transfection in some experiments.

HEK293T or NPC43 cells were pre-inoculated in 24-well plates and co-transfected with the luciferase reporter plasmid and the corresponding plasmids using PEI as described above. After 30-40 h of transfection, cell lysates were analyzed using the Dual-Luciferase® Reporter Assay System (Promega) following the manufacturer’s instructions. The pRLTK (Renilla) luciferase reporter was used as a control. A GloMAX™ 96 microplate luminometer (Promega) was used to measure the bioluminescence.

### Statistical analysis

The data shown are the mean ± SEM of at least three independent experiments. Paired Student’s *t*-test analysis of variance was used for statistical analysis unless otherwise stated. *p*<0.05 indicates statistical significance.

## Results

### BRRF1 promotes expression of BTN2A1 and BTN3A1 in NPC43 cells

Based on our previous report, we found that BRRF1 may be associated with the expression of BTN2A1 and BTN3A1 in NPC cells [9]. To investigate the possibility, we first overexpressed BRRF1 in NPC cell lines, HK1 and HONE1, which do not harbour EBV (i.e. EBV-negative) and EBV-positive cell lines (C666-1 and NPC43), and detected the gene expression of *BTN2A1, BTN3A1*, and *NLRC5* by RT-qPCR. The results showed no difference in the expression of these genes in HK1, HONE1, or C666-1 cell lines (**Figure S1**). However, in the most clinically relevant NPC43 cells, the expression of *BTN2A1* and *BTN3A1* was increased (**Figure 1A**). This was confirmed by Western blot analysis where BRRF1 indeed upregulated BTN2A1 and BTN3A1 protein expression in NPC43 cells (**Figure 1B**). NLRC5 was not affected by BRRF1, suggesting a different mechanism to LMP1 as we reported previously [9].

**Figure 1.**
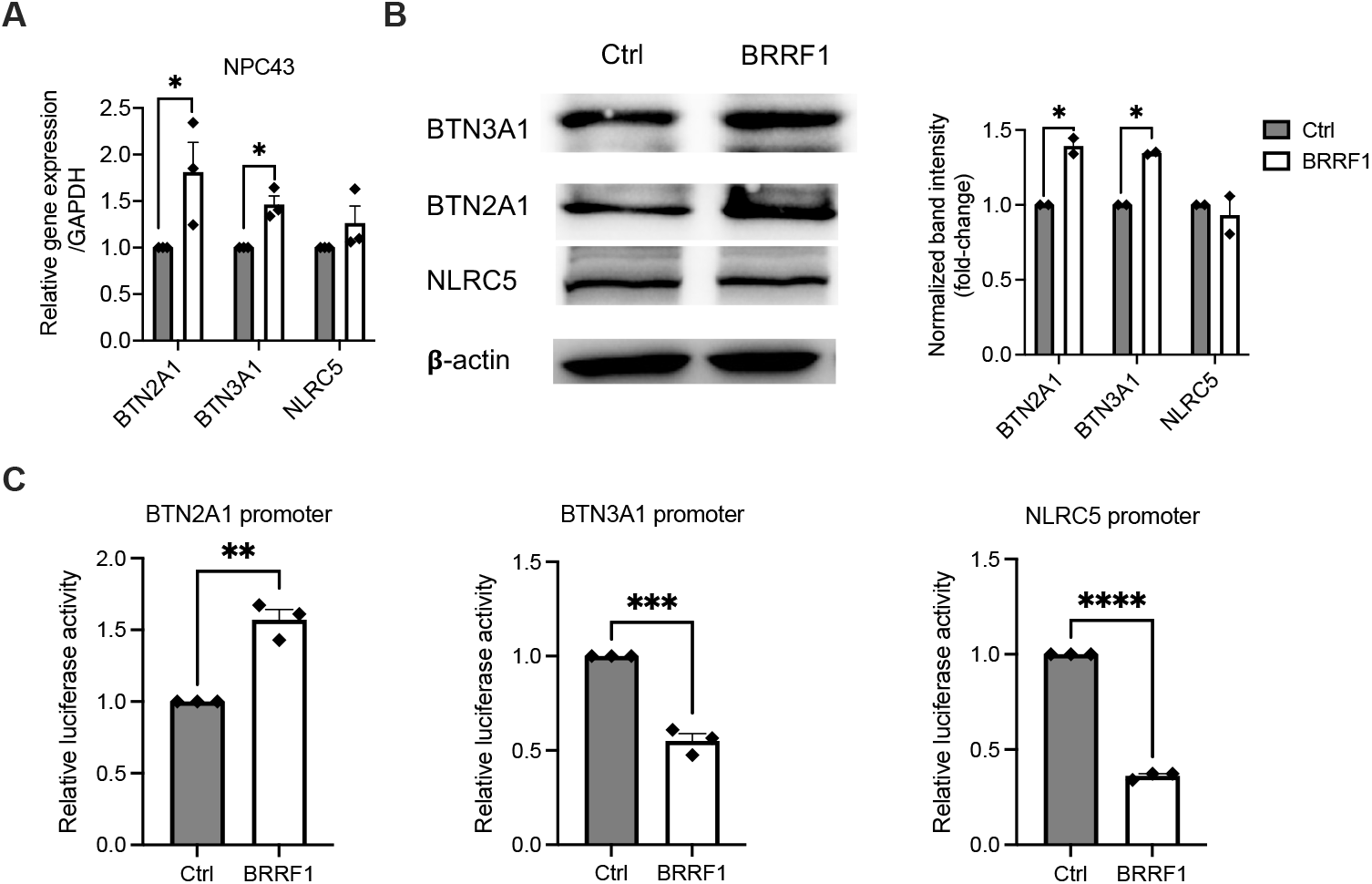
BRRF1 overexpression induced protein expression of BTN2A1 and BTN3A1 in EBV^+^ NPC cells. (A) NPC43 cells were transfected with BRRF1 and assessed for *BTN3A1, BTN2A1*, and *NLRC5* gene expression by RT-qPCR. (B) Western blot analysis of BTN3A1, BTN2A1 and NLRC5 protein levels of BRRF1 overexpression in NPC43 cells 24 h compared to β-actin. Representative immunoblots and the band intensity normalized to the control as column graphs are shown. (C) Luciferase activities from promoters for BTN2A1, BTN3A1, and NLRC5 in 293T cells transfected with plasmids encoding BRRF1 or vector control for 1 day. Data shown as mean±SEM from ≥3 independent experiments. Student’s *t*-test was performed for statistical analysis. \**P* < 0.05, \*\**P* < 0.01, \*\*\**P* < 0.001.

Next, to test whether BRRF1 could directly trigger the transcription of these genes, the luciferase reporter assay was performed in HEK293T cells using plasmids encoding the promoters of BTN2A1, BTN3A1, and NLRC5. Interestingly, BRRF1 triggered the BTN2A1 promoter activity but not BTN3A1 and NLRC5 (**Figure 1C**). Based on these results, we demonstrated that BRRF1 can stimulate expression of BTN2A1 in NPC43 cells, which has not been shown before.

### Analysis of transcriptome following BRRF1 expression in NPC43 cells

To investigate the mechanism of BRRF1 induction of BTN2A1, we overexpressed BRRF1 in NPC43 cells and performed total RNA sequencing. Among the top 50 differentially expressed genes, we found significantly 19 upregulated genes and 31 downregulated genes compared to control (**Figure 2A**). Some of these genes appear to be related to Ras GTPase binding and signaling, extracellular structure, and MAPK or PI3K-Akt signaling pathways based on KEGG and GO enrichment analyses (**Figure 2B and 2C**). These genes will serve as the basis to investigate the mechanism of BRRF1 function related to BTN2A1.

**Figure 2.**
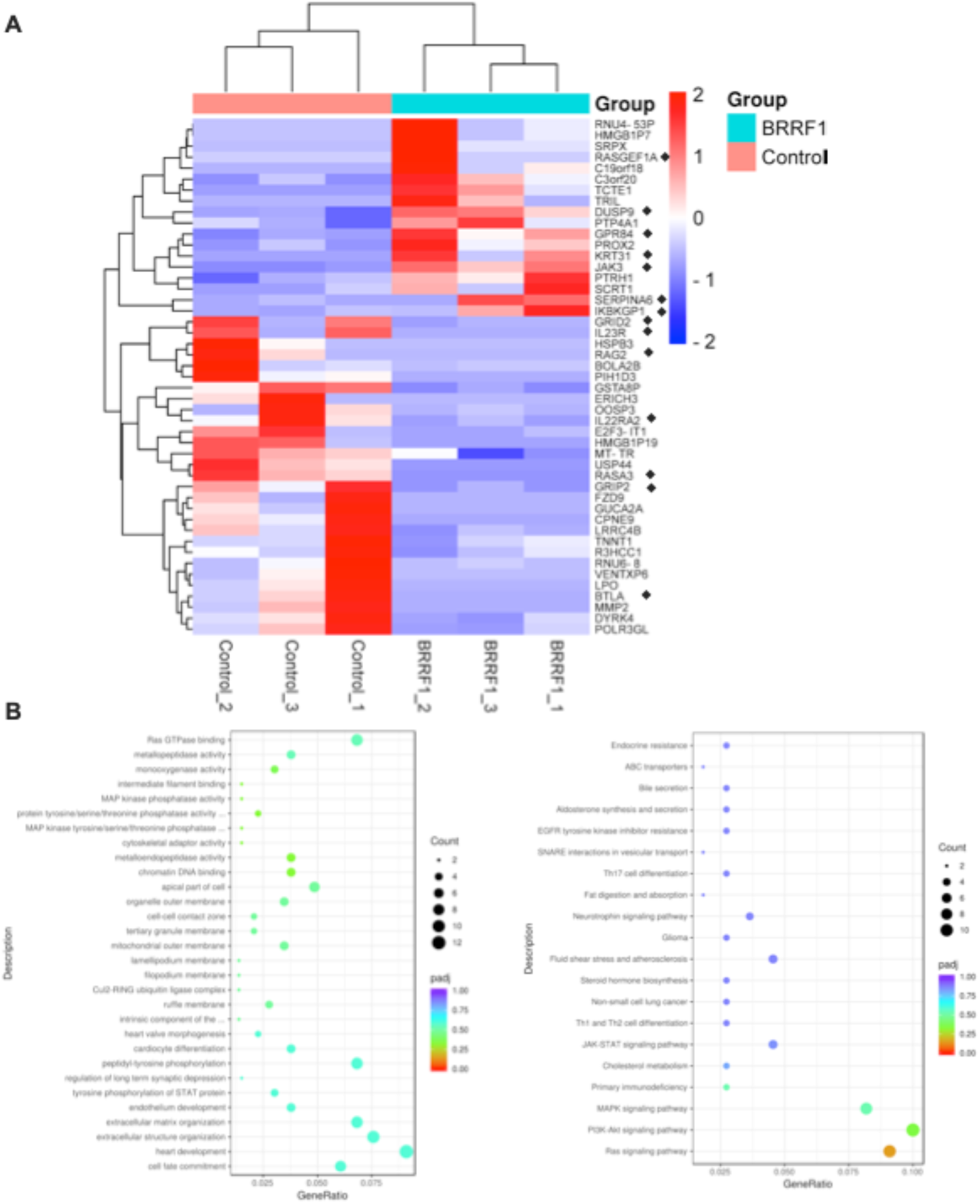
RNA sequencing analysis of genes altered by BRRF1 overexpression in NPC43 cells. (A) Heatmap of the RNA sequencing data in control and BRRF1 overexpression groups. Three samples from each group were used for the clustering. The degree of gene expression of BRRF1 compared to control ranged from blue (downregulated) to red (upregulated) in a log2 scale. (B) KEGG and (C) GO analysis of the differentially expressed genes in BRRF1 versus control.

We selected genes of interest based on the available information of their functions that may be related to transcriptional control or signaling pathways. 11 up-regulated genes (**Table S3**), and 9 down-regulated genes (**Table S4**) were chosen for RT-qPCR verification. Indeed, overexpression of the BRRF1 in NPC43 upregulated the expression of *JAK3* and *RhoB* genes while downregulating the *IL22RA2* gene (**Figure 3A**). The other genes showed no statistically significant differences (**Figure S2** and **Figure S3**). By Western blotting, overexpression of BRRF1 in NPC43 cells after 24 h led to increased JAK3 and decreased IL-22RA2 protein expression, with no effect on RhoB (Figure 3B). Antibody against Myc was used to detect BRRF1 expression in the cells.

**Figure 3.**
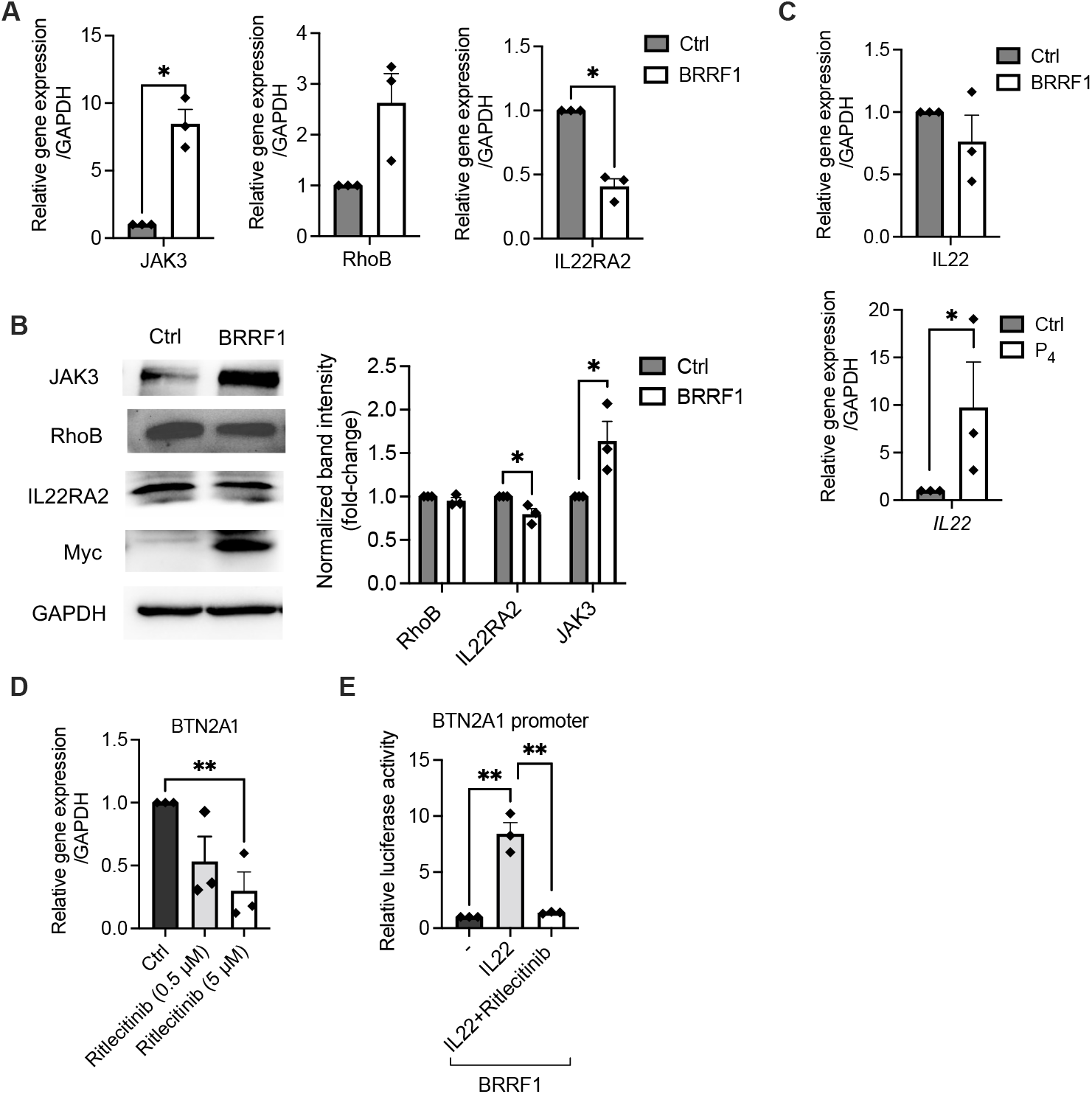
BRRF1 induces BTN2A1 expression through the IL22 and JAK3 pathways. (A) NPC43 cells were transfected with BRRF1 for 3 h and assessed for *JAK3, RhoB*, and *IL22RA2* gene expression by RT-qPCR. (B) Representative immunoblots from Western blot analysis of BRRF1(Myc), RhoB, IL22RA2, and JAK3 expression following transfection of NPC43 cells by BRRF1 with 24 h. Band intensities are calculated compared to the control. (C) NPC43 cells were transfected with BRRF1 or treated with P_4_ and assessed for *IL22* gene expression by RT-qPCR. (D) RT-qPCR of *BTN2A1* gene expression of NPC43 cells treated with JAK3 inhibitor ritlecitimib (0.5 and 5 μM) or DMSO control. (E) Luciferase reporter assays using BTN2A1 promoter in NPC43 cells following BRRF1 overexpression with or without IL-22 (10 ng/ml) treatment. Column graphs represent data as mean±SEM from ≥3 independent experiments. Student’s *t*-test was performed for statistical analysis. \**P* < 0.05, \*\**P* < 0.01.

IL-22RA2 (or IL-22 binding protein, IL-22BP) is a naturally secreted monomeric protein whose primary function is to block the interaction of IL-22 with IL-22R by binding specifically to extracellular IL-22 and, thereby, inhibit downstream responses [25]. IL-22, IL-22R, and IL-22RA2 interactions play an important role in health and diseases and are involved in regulating cancer and homeostasis [26]. Therefore, the mRNA and protein downregulation of IL22RA2 in NPC43 cells (**Figure 3A** and **3B**) caused by BRRF1 will likely promote IL22 binding with IL22R. Besides the decreased expression of IL22RA2, we wondered if BRRF1 can stimulate the expression of IL22. However, RT-qPCR analysis showed that BRRF1 did not directly induce gene expression of IL22, but stimulation of NPC43 cells using P_4_ (10 mM) could do so (**Figure 3C**). This suggests that EBV reactivation can induce the expression of IL22 and may be caused by EBV genes other than BRRF1. Moreover, BRRF1 did not influence RhoB protein expression (**Figure 3A and B**), indicating that the Ras-GTPase pathway is not a major target by BRRF1.

### BRRF1 induces BTN2A1 expression via the JAK3-STAT3 pathway

In consistence to RNA-seq and RT-qPCR data, BRRF1 increased JAK3 protein levels (**Figure 3B**), suggesting that the JAK signaling cascade pathway could be activated by BRRF1. To test the other genes belonging to this family, BRRF1 also induced higher mRNA expression of *JAK1* and *TYK2* but not *JAK2* (**Figure S4**). Further, using an inhibitor against JAK3, ritlecitinib, significantly decreased the gene expression level of *BTN2A1* in NPC43 cells (**Figure 3D**). In addition, IL-22 synergistically upregulated the *BTN2A1* promoter activity with BRRF1, and this effect was abrogated when the JAK3 was inhibited (**Figure 3E**). On the protein level, BRRF1 transfection in NPC43 cells lead to increased JAK3 and p-JAK3 expression, but not TYK2/p-TYK2, JAK1/p-JAK1, and JAK2/p-JAK2, which confirmed that the JAK3 pathway play an important role for BRRF1 to induce BTN2A1 expression in NPC43 cells (**Figure 4A** and **Figure S5A**). The use of JAK3 inhibitor corroborated with this finding. Ritlecitinib treated NPC43 cells can decrease JAK3/p-JAK3, and successfully downregulated the effect of BRRF1 induction of BTN2A1 protein levels. No significant effects were shown for the other JAK family proteins. Interestingly, JAK3 inhibition appears unable to alter the expression of BTN3A1, indicating that BRRF1 induction of BTN3A1 may be dependent on other pathways.

**Figure 4.**
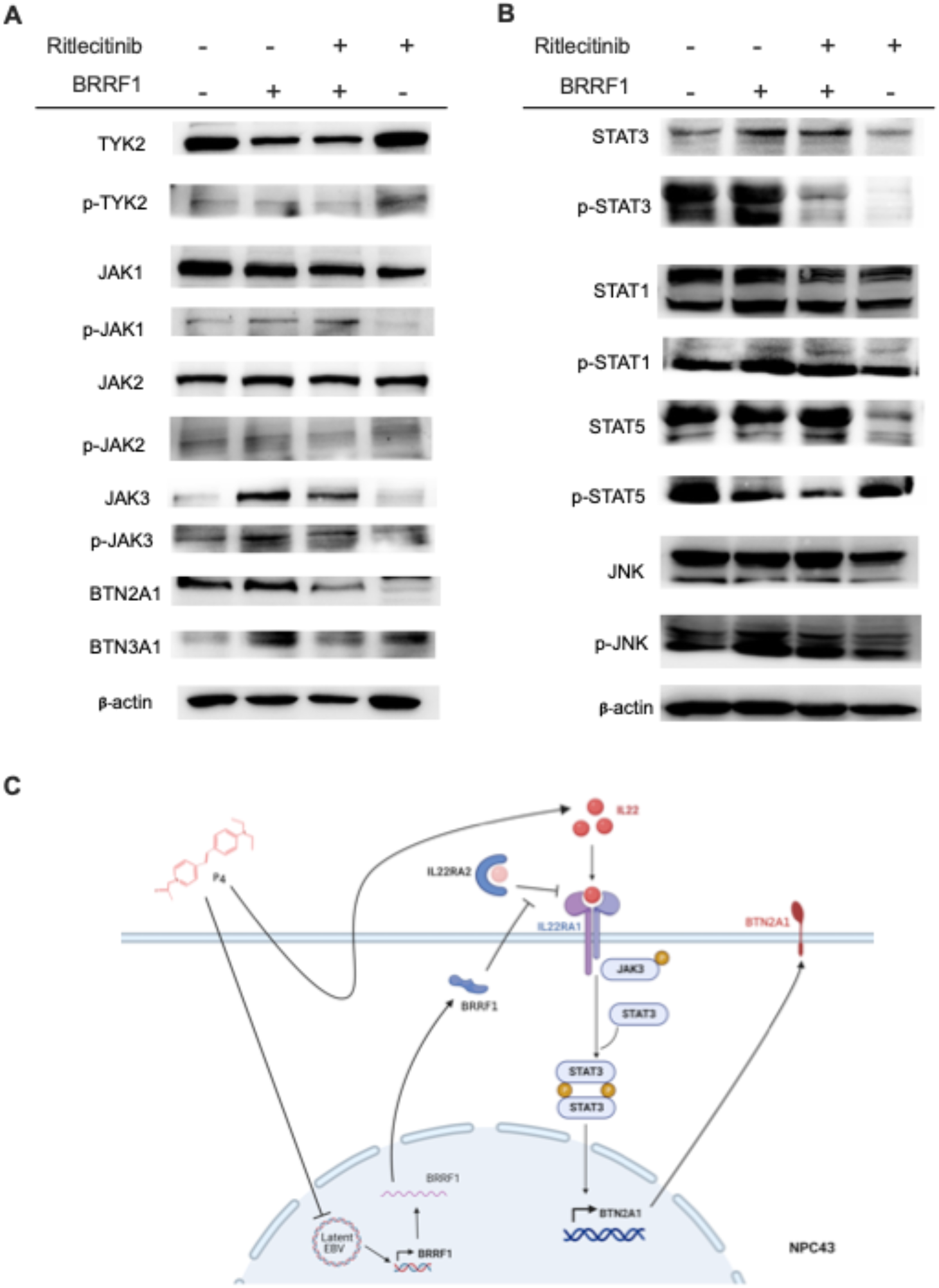
BRRF1 induces the BTN2A1 protein level through the JAK3-STAT3 pathway. (A) Western blot analysis of TYK2, p-TYK2, JAK, p-JAK1, JAK2, p-JAK2, JAK3, p-JAK3, BTN2A1, and BTN3A1, as well as (B) STAT3, p-STAT3, STAT1, p-STAT1, STAT5, p-STAT5, JNK, and p-JNK protein expression of NPC43 cells by BRRF1 overexpression with or without ritlecitinib (5 μM) treatment. Representative immunoblots are shown. (C) Schematic diagram of the role of BRRFI, and P_4_/IL-22 in the induction of BTN2A1 expression via JAK3-STAT3 pathway.

Upon binding to IL-22R1, IL-22 signals via the JAK-STAT pathways primarily activate STAT1, STAT3, and STAT5 [27]. To further examine the effect of BRRF1 in cooperating with IL-22, we measured the protein expression of STAT3, STAT1, and STAT5 in NPC43 cells under different treatments. As shown in **Figure 4B**, BRRF1 induced STAT3 and p-STAT3 protein levels, but not STAT1/p-STAT1, STAT5/p-STAT5 or JNK/p-JNK. Inhibitor against JAK3 abrogated this effect, which indicates that JAK3 was responsible for activating STAT3 for downstream induction of BTN2A1 (**Figure 4B and Figure S5B**).

## Discussion

During latency, the gene expression by EBV remains subtle and viral persistence is maintained by latent proteins such as EBNA1 and LMP1, which could also lead to tumor transformation of the host cells [28]. Though EBV genes towards oncogenesis is well-characterized, there is also an opportunity to study their role (if any) in promoting anti-tumor immune cell functions. The difficulty for current immunotherapy against NPC is to overcome the immunosuppressive tumor microenvironment. The use of immune checkpoint inhibitors or to enhance anti-tumor T cell functions have shown limited success [29], therefore, other avenues can be explored to resolve the underlying reasons. Along this line, studying the existing EBV genes within the NPC tumor cells could augment cell-based immunotherapy. Here, we demonstrated that the EBV gene BRRF1, which can be triggered by an EBV-targeting probe, upregulates the expression of BTN2A1 that would support Vγ9Vδ2 T cell adoptive immunotherapy against NPC. BRRF1 is classified as a lytic gene that encodes a 34 kDa protein and can be activated by the transactivator BZLF1 during EBV replication [30]. The protein has been found in both the nucleus and cytoplasm [30, 31]. BRRF1 has also been found to activate the BZLF1 promoter and its own expression [31, 32]. Further, it was shown that BRRF1 can activate promoters that contain p53, AP-1, CREB and NF-κB response elements [33]. Therefore, BRRF1 has the possibility to mediate BTN expression as shown in our data. BRRF1 appears to directly trigger BTN2A1 promoter activity and lead to the increased expression of BTN2A1 protein. In concordance with IL-22, BRRF1 can downregulate the inhibitory IL-22 binding protein, and likely amplify the expression of BTN2A1 in a JAK3-STAT3 pathway dependent manner. These functions have not been attributed for BRRF1 before.

BTN2A1 is one of the important molecules in activating and ensuring the function of Vγ9Vδ2 T cells against tumor cells. BTN2A1 and BTN3A1 forms a complex associated with phosphoantigen to be recognized by the Vγ9Vδ2-TCR that drives the downstream signaling for the cells [18]. In the NPC tumor microenvironment, the expression of these molecules is lacking [9], and this would render γδ T cells adoptive therapy ineffective when they reach the tumor site. The discovery from this study that BRRF1 can induce BTN2A1 will enhance the tumoricidal acitivity of Vγ9Vδ2 T cells, as we showed previously [9]. As far as we know, the induction pathway of BTN2A1 remains elusive, and our data suggests that BRRF1, JAK3 and STAT3 could be possible candidates. Therefore, BRRF1 may be used as a tool to modify the tumor prior to Vγ9Vδ2 T cell immunotherapy against NPC. In addition, the finding that IL-22 could stimulate BTN2A1 is also surprising. IL-22 is part of the IL-10 family of cytokines and can target non-hematopoietic cells, including epithelial cells and fibroblasts, and can affect tissues in the gut, lung, liver, pancreas, breast, skin, kidney, and thymus [34]. It can be produced by cells of the lymphoid lineage, including αβ CD4^+^ and CD8^+^ T cells and γδ T cells (Interleukin (IL)-22 and IL-17 are co-expressed by Th17 cells and cooperatively enhance expression of antimicrobial peptides [35-39]. However, the role of IL-22 in promoting or suppressing tumor remains controversial. In colon cancer, IL-22 could activate STAT3 and cause progression [40], but there is no association of IL-22 being the causative effect on tumor transformation [41]. On another note, IL-22BP has a regulatory role as it is constitutively expressed in various tissues, but in cancer, it appears to be decreased that can allow IL-22-induced tumor growth [42]. Of note, IL-22BP is constitutively expressed in various tissues, and it counteracts the inflammatory properties of IL-22 [43]. However, in cancer, IL-22BP were found to be decreased, which would then allow IL-22 effects [44]. However, results from this study showed that IL-22 in conjunction with BRRF1 could stimulate the expression of BTN2A1, with the downregulation of IL-22RA2 by BRRF1. This is an interesting observation as NPC43 cells could express IL-22 following induced EBV-reactivation, but the amount produced is most likely less than lymphoid cells and likely can assist anti-tumor Vγ9Vδ2 T cells. However, when IL-22 producing Th17, NKT, Th22 cells infiltrate into the tumor, the high amount of IL-22 can offset this and eventually lead to tumor progression [35, 45, 46]. In this consideration, our data suggests that BRRF1-induced downregulation of IL-22RA2 could help IL-22-induced BTN2A1 but might also allow IL-22-induced tumor progression if IL-22 levels are high. This should be considered in future development of Vγ9Vδ2 T cell therapy in combination with BRRF1 in patients. In summary, we uncovered a new mechanism to induce BTN2A1 in NPC by BRRF1 and/or IL-22, with the downregulation of IL-22BP, and activation of the JAK3-STAT3 pathways (**Figure 4C**). BRRF1, as well as LMP1 [9], may have uses in transducing other tumors for promoting BTN2A1 and BTN3A1 expression prior to the adoptive transfer of Vγ9Vδ2 T cells; this may be one path of enhancing the immunotherapy. Thus, the new findings here can provide insights to exploit tumor-associated viral genes in various cancers for immunotherapy.

## Acknowledgements

The authors thank Ms Yee Man Lee for laboratory and experimental support and contribution to this work.

## Data availability

All relevant data can be found within the article and its supplementary information. RNA-seq data can be made available upon request.

## Figure legends

**Figure S1. BTN2A1, BTN3A1, and NLRC5 gene expression were measured in different NPC cell lines following BRRF1 overexpression**. (A) HK1, (B) HONE1, and (C) C666-1 cell lines were transfected with BRRF1 in 3 h and assessed for *BTN3A1, BTN2A1*, and *NLRC5* gene expression by RT-qPCR.

**Figure S2. Up-regulated gene mRNA expression was measured in Ctrl and BRRF1 overexpression NPC43 cells by RT-qPCR**. *ADAMTS9, RASGEF1A, KRT31, SEPRINA6, AKR1D1, GRP84, RASGRF1, IKBKG1*, and *DUSP9* gene mRNA expression was measured by RT-qPCR.

**Figure S3. Down-regulated gene mRNA expression was measured in CTRL and BRRF1 overexpression NPC43 cells by RT-qPCR**. *RASA3, IL23R, RAG2, GRIP2, GRID2, MAP3KTCL, ABCB11, BTLA* gene mRNA expression was measured by RT-qPCR.

**Figure S4. NPC43 cells were transfected with BRRF1 for 3 h and assessed for *JAK1, JAK2*, and *TYK2* gene expression by RT-qPCR**.

**Figure S5. Western blot analysis of different JAK and STAT pathway protein expression of NPC43 cells with different treatment groups measured by band intensities**. (A) JAK family proteins, BTN2A1 and BTN3A1, and (B) STAT family proteins relative band intensities normalized to β-actin are shown. Data from 3 independent experiments are shown as mean ± SEM in the column graphs. Student’s *t*-test was performed. \**P* < 0.05, \*\**P* < 0.01, \*\*\**P* < 0.001, \*\*\*\**P* < 0.0001.

